# Seed hemicelluloses tailor mucilage properties and salt tolerance

**DOI:** 10.1101/2020.08.03.234708

**Authors:** Bo Yang, Florian Hofmann, Björn Usadel, Cătălin Voiniciuc

## Abstract

- While Arabidopsis seed coat epidermal cells have become an excellent genetic system to study the biosynthesis and structural roles of various cell wall polymers, the physiological function of the secreted mucilaginous polysaccharides remains ambiguous. Seed mucilage is shaped by two distinct classes of highly substituted hemicelluloses along with cellulose and structural proteins, but their interplay has not been explored.
- We deciphered the functions of four distinct classes of cell wall polymers by generating a series of double mutants with defects in heteromannan, xylan, cellulose, or the arabinogalactan protein SALT-OVERLY SENSITIVE 5 (SOS5), and evaluating their impact on mucilage architecture and on seed germination during salt stress.
- We discovered that *muci10* seeds, lacking heteromannan branches, had elevated tolerance to salt stress, while heteromannan elongation mutants exhibited reduced germination in CaCl_2_. In contrast, xylan made by *MUCILAGE-RELATED21* (*MUCI21*) was found to be required for the adherence of mucilage pectin to microfibrils made by CELLULOSE SYNTHASE5 (CESA5) as well as to a SOS5-mediated network.
- Our results indicate that the substitution of xylan and glucomannan in seeds can fine-tune mucilage adherence and salt tolerance, respectively. The study of germinating seeds can thus provide insights into the synthesis, modification and function of complex glycans.

## Introduction

Cellulose microfibrils are deposited around plant cells and enmeshed in a complex matrix of hemicelluloses, pectin, and, to a lesser extent, structural proteins. The roles of specific classes of cell wall polymers have been difficult to study even in model organisms. For instance, *Arabidopsis thaliana* has nine *CELLULOSE SYNTHASE-LIKE A (CSLA)* genes that are at least putatively involved in the synthesis of heteromannan (HM), a class of hemicellulose mainly built of β-1,4-linked mannosyl units. While HM polymers could store carbon to feed growing seedlings or directly control cell wall structure (Schröder *et al.*, 2009), their physiological roles in Arabidopsis are poorly understood. Genetic disruption of *CSLA7* is embryo-lethal, but *csla2 csla3 csla9* triple mutant stems had no phenotypic changes despite lacking detectable HM (Goubet *et al.*, 2009). Significant insights into the biosynthesis and functions of various cell wall components, including HM, have been gained using the Arabidopsis seed coat as a genetic model (Šola *et al.*, 2019). The seed coat epidermis secretes large amounts of polysaccharides that rapidly swell upon hydration to release non-adherent mucilage as well as an adherent capsule. Unbranched pectin is the dominant mucilage component, but the adherent capsule also contains hemicellulosic polymers typical of secondary walls (Voiniciuc *et al.*, 2015c), which are deposited after cells expand.

In the past decade, several classes of carbohydrate-active enzymes have been found to influence mucilage content and properties (Griffiths & North, 2017; Šola *et al.*, 2019). At least three genes are required to maintain pectin adherence to the seed surface (Fig. 1a): *CELLULOSE SYNTHASE (CESA5), SALT-OVERLY SENSITIVE5 (SOS5)* and *MUCILAGE-RELATED21/MUCILAGE-MODIFIED5 (MUCI21/MUM5).* CESA5 is a member of the cellulose synthesis complex (Sullivan *et al.*, 2011; Mendu *et al.*, 2011; Harpaz-Saad *et al.*, 2011; Griffiths *et al.*, 2015), while the SOS5 arabinogalactan protein could be part of a mucilage proteo-glycan or a kinase signalling pathway (Harpaz-Saad *et al.*, 2011; Griffiths *et al.*, 2014; Basu *et al.*, 2016). Although its predicted xylosyltransferase activity remains to be confirmed *in vitro* (Voiniciuc *et al.*, 2015a; Zhong *et al.*, 2018), MUCI21 is required to substitute xylan with xylose branches (Voiniciuc *et al.*, 2015a) that facilitate pectin-cellulose interactions (Ralet *et al.*, 2016). Galactoglucomannan, another branched hemicellulose in Arabidopsis mucilage, is elongated by CSLA enzymes and substituted by MANNAN α-GALACTOSYLTRANSFERASE1/MUCILAGE-RELATED10 (MAGT1/MUCI10; Yu *et al.*, 2014, 2018; Voiniciuc *et al.*, 2015b). Unlike xylan, branched HM maintains cellulose deposition and pectin density without appearing to influence mucilage adherence (Fig. 1a).

**Fig. 1.**
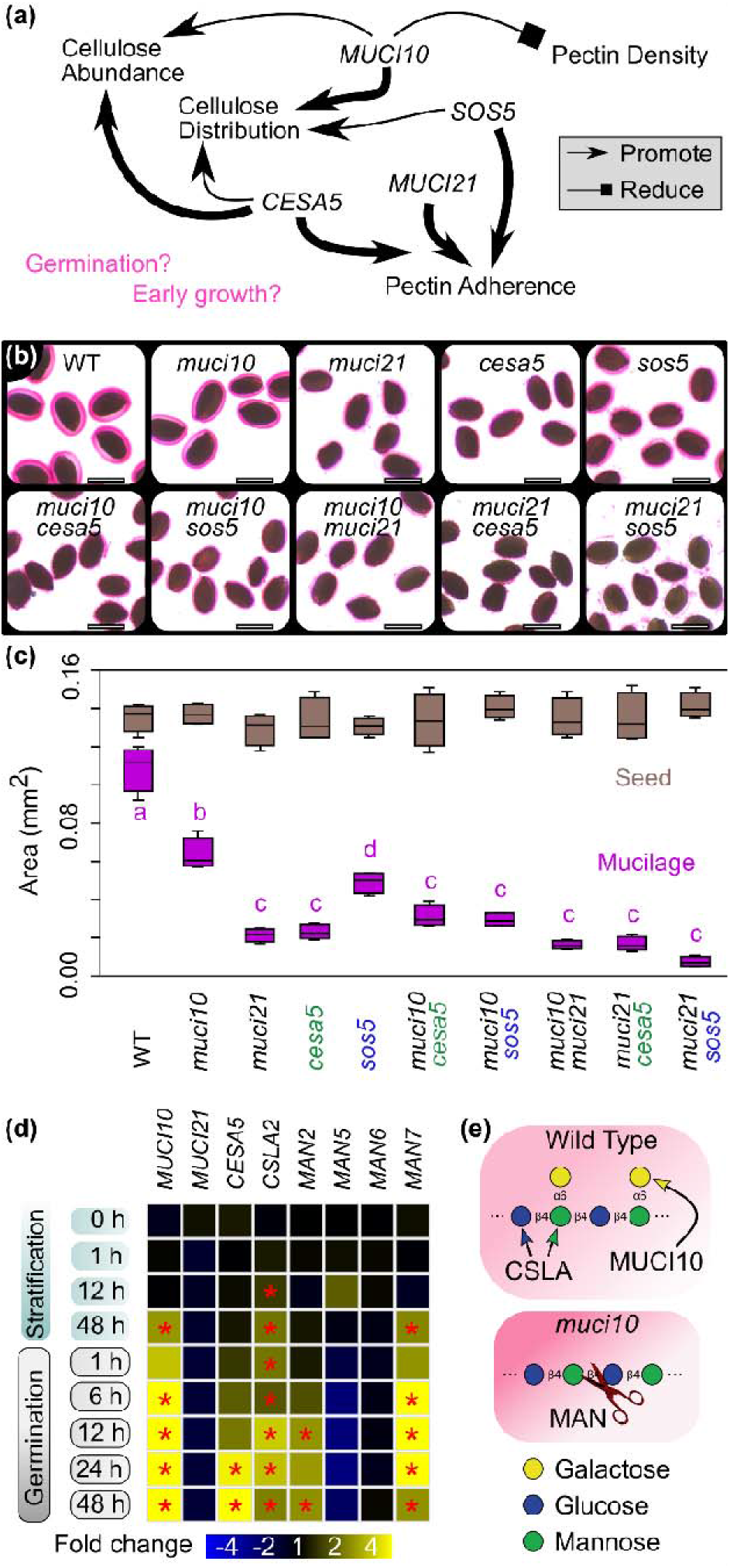
Impact of different players on mucilage properties. (a) Schematic of previously reported functions of four genes on Arabidopsis seed mucilage properties. Genetic interactions between these players and their physiological roles remain unknown. (b) Wild-type (WT) and mutant seeds were gently mixed in water and RR was used to stain adherent pectin (pink). Scale bars = 0.6 mm. (c) Box plots of projected seed and mucilage areas of four biological replicates (each with around 17 seeds) per genotype. Letters denote significant differences (one-way ANOVA with Tukey test, P < 0.01). (d) Transcriptional changes (asterisks; P < 0.001) during stratification and germination relative to dry seeds (0 h), profiled in GENEVESTIGATOR. (e) Schematic of HM structure in wild-type and *muci10* mutant, which is expected to be more accessible to cleavage or re-modelling by β-1,4-mannanases (MAN).

Biochemical and histological analyses of double mutants have clarified how SOS5 and cellulosic ray-like structures provide two distinct mechanisms to anchor pectin to seeds (Griffiths *et al.*, 2014, 2016; Ben Tov *et al.*, 2018). The contrasting roles of the two hemicelluloses on mucilage properties have yet to be evaluated in detail. The physiological roles of Arabidopsis seed mucilage are still ambiguous, even though angiosperm seed coats have been involved in seed dormancy, dispersal and germination (Western, 2012; North *et al.*, 2014). In contrast to the Columbia wild type, Arabidopsis varieties with impaired mucilage release (Saez-Aguayo *et al.*, 2014) or adherence (Voiniciuc *et al.*, 2015a) have elevated buoyancy and could be dispersed on water. Seed germination is essential for plant establishment and is highly sensitive to salt stress. In this study, we therefore explored how genes affecting different wall polymers modulate mucilage properties, seed germination and early growth under salt stress (Fig. 1a).

## Materials and Methods

### Plant materials

Mutations were genotyped using primers listed in Table S1 and Touch-and-Go PCR (Berendzen *et al.*, 2005). The double mutants generated in this study are available from the Nottingham Arabidopsis Stock Center. Plants were grown in climate-controlled chambers as previously described (Voiniciuc *et al.*, 2015b). The germination assays were performed using seeds produced by plants grown individually in 8 cm round pots at 100-120 μE m^−2^ s^−1^ light, 22°C and around 60% relative humidity. Flowering plants were staked and mature, dry seeds (~10 weeks) were harvested, separated from the chaff and stored in separate paper bags (one per plant) in a temperature-controlled laboratory (~23°C, 40 to 50% humidity).

### Microscopic analyses

Seeds were stained with 0.01% ruthenium red (RR) in 24-well plates and quantified in Fiji (https://fiji.sc/; Schindelin et al., 2012) using established protocols (Voiniciuc *et al.*, 2015b). For staining without shaking, seeds were imbibed in 300 μL of 0.01% RR solution for 15 min. Images were acquired with two stereomicroscope-camera setups: MZ12 with DFC 295, or M165FC with MC170 HD (all from Leica). Mucilage immunolabeling with CCRC-M139 (Carbosource, Complex Carbohydrate Research Center) and counter-staining with S4B (Direct Red 23; Sigma Aldrich) was performed using a published protocol and Leica TCS SP8 confocal setup (Voiniciuc, 2017). Germinated seeds were stained with calcofluor white and propidium iodide (0.05%, w/v, for both dyes) for 10 min, rinsed well with water, and imaged on a Zeiss Imager.Z2 with a 10x Plan-Fluar (NA 0.30), Axiocam 506, and DAPI/Texas Red filters.

### Biochemical analyses

Total mucilage was extracted with a ball mill, hydrolyzed, and quantified via high performance anion exchange chromatography with pulsed amperometric detection (HPAEC-PAD) as previously described (Voiniciuc & Günl, 2016). The quantification of mucilage detachment via HPAEC-PAD has also been described in detail (Voiniciuc, 2016). HPAEC-PAD of mucilage was conducted on a Dionex system equipped with CarboPac PA20 columns (Voiniciuc & Günl, 2016). For alcohol-insoluble residue (AIR) isolation, all material (72 h post-stratification) from four biological replicates was pooled, finely ground and sequentially washed with 70% ethanol, chloroform:methanol (1:1, v/v) and acetone. Monosaccharide content of germinated seed AIR after 2 M trifluoroacetic acid hydrolysis was analyzed on a Metrohm 940 Professional IC Vario (Voiniciuc *et al.*, 2019), equipped with Metrosep Carb 2-250/4.0 guard and analytical columns.

### Seed germination assay

All germination assays were performed in sterile 24-well culture plates (VWR International; 734-2779), using 500 μL of the specified solution and dry seeds (typically 20, but up to ~100 worked) from a single plant per well. The four corners had only water and the plates were sealed with lids and 3M micropore tape to reduce desiccation. Replicates from high-quality seed lots were distributed to avoid positional bias, and at least three biological replicates per genotype showed consistent results. Seeds were hydrated in 500 μL of distilled water, 150 mM CaCl2 or 150 mM NaCl directly in the plate, or first de-mucilaged via ball mill extraction in water (Voiniciuc & Günl, 2016) before rinsing and being transferred in the final solvent (500 μL) to the plates. Floating seeds were counted as the number remaining in the center of each well, atop the solution. Plates were stratified for 66 h (dark, 4°C), transferred to a phytochamber (22°C, 100 μE m^−2^ s^−1^ constant light), and then imaged every 24 h with a Leica M165FC stereomicroscope. Seeds were defined as germinated if radicle length was >70 μm, when quantified in Fiji (line tool).

To compare ionic and osmotic effects, germination assays were performed in 150 mM CaCl_2_ or MgCl_2_ salts, 450 mM sorbitol, and 61 mM polyethelene glycol (PEG) 4000, all with an equal osmotic pressure (1.11 MPa) based on the van ’t Hoff formula and experimental data (Money, 1989). Radicle protrusion versus elongation effects were tested by switching water and 150 mM CaCl_2_ at 24 h post-stratification following three sequential 450 μL solvent exchanges.

### Figures and statistical analysis

Micrographs were processed uniformly in Fiji. Numerical data were plotted as bar graphs in Microsoft Excel 365 or as box/violin/jitter plots in the Past 4 statistics software package (https://folk.uio.no/ohammer/past/; Hammer *et al.*, 2001)). Panels were assembled in Inkscape (https://inkscape.org/). ATH1 microarray expression, including GSE20223 dataset (Narsai *et al.*, 2011), was visualized in GENEVESTIGATOR Professional (https://genevestigator.com/). Two-samples and multiple samples statistics were performed in Excel and Past 4, respectively. Carbohydrates were drawn according to the Symbol Nomenclature for Glycans (SNFG).

## Results and Discussion

### Mucilage adherence requires multiple wall polymers, except HM

To dissect the roles of the four genes listed in Fig. 1a, we generated a series of double mutants with defects in HM, xylan, cellulose or an AGP (SOS5). We crossed the *muci10-1* (Voiniciuc *et al.*, 2015b) and *muci21-1* (Voiniciuc *et al.*, 2015a) hemicellulose mutants to each other, as well as to *cesa5-1* (Mendu *et al.*, 2011) and *sos5-2* (Harpaz-Saad *et al.*, 2011). After shaking and RR staining, the seeds of all single and double mutant combinations had wild-type seed area but were surrounded by smaller mucilage capsules (Fig. 1b,c). *MUCI21, CESA5* or *SOS5* were epistatic to *MUCI10* in terms of adherent mucilage size. While all mutants produced wild-type percentages of rhamnose and galacturonic acid in total mucilage extracts (Table S2), significant reductions in minor sugars were associated with *muci10* (galactose and mannose) and *muci21* (xylose) mutations (Fig. 2a). Consistent with previous results (Griffiths *et al.*, 2014), *cesa5* and *sos5* mutations did not alter matrix polysaccharide composition. The *muci10 muci21* double mutant phenocopied the biochemical deficiencies of the respective single mutants, indicating that xylan and HM substitution can be uncoupled in the seed coat.

**Fig. 2.**
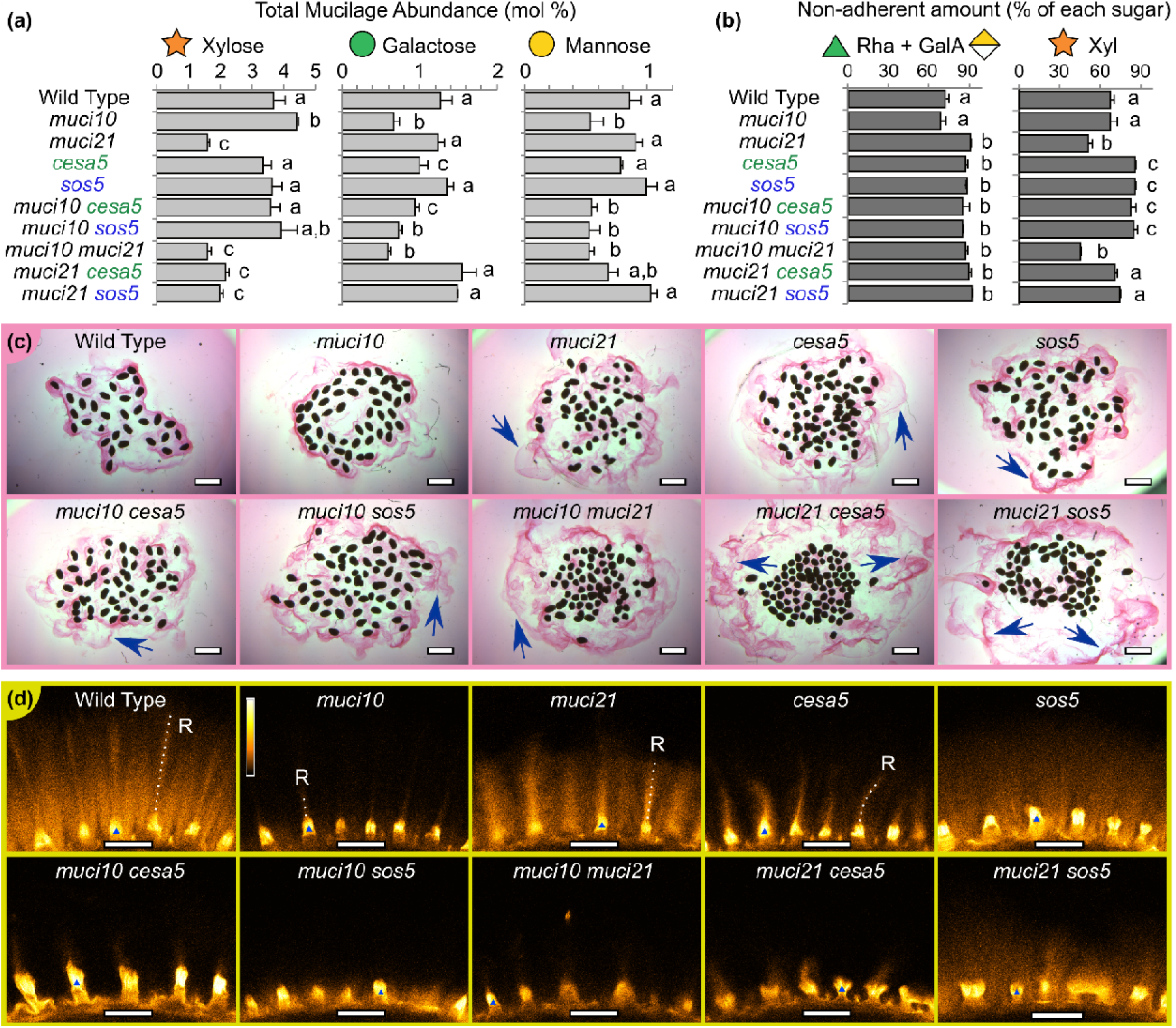
Mucilage polysaccharide composition and distribution. (a) Relative abundance of hemicellulose-derived monosaccharides in total mucilage. (b) The non-adherent proportion of mucilage pectin (sum of rhamnose and galacturonic acid) and xylan (built of xylose residues). Data show mean + SD of four biological replicates, except only two for *sos5* in (b), and letters denote significant differences (one-way ANOVA with Tukey test, P < 0.05). (c) Hydration of seeds in RR solution, without shaking. Blue arrows indicate non-adherent mucilage. (d) S4B staining of cellulose, coloured using Orange Hot LUT in Fiji (see bar in *muci10* subpanel). Blue triangles mark volcano-shaped columellae on the seed surface, and the dashed lines indicate cellulosic rays (labelled R). Scale bars = 1 mm (c), 50 μm (d).

Sequential mucilage extractions (Fig. 2b and Table S3) as well as direct hydration in RR solution (Fig. 2c) showed that more pectin detached from seeds containing *muci21, cesa5,* and/or *sos5* mutations compared to wild-type and *muci10*. Xylan detachment increased proportional to that of pectin in mutants lacking *CESA5* and/or *SOS5* (Fig. 2b; Table S3), consistent with covalent linkages between these polymers (Ralet *et al.*, 2016; Voiniciuc *et al.*, 2018). Unbranched xylan epitopes, labelled by the CCRC-M139 monoclonal antibody (Ruprecht *et al.*, 2017), closely surrounded *muci21* and *cesa5* seeds, but were further from the surface of *sos5* and other genotypes (Fig. S1a,b), proportional to the RR-stained adherent capsule size (Fig. 1b,c).

Each mutation also had distinct effects on S4B staining, which primarily detects cellulose (Anderson *et al.*, 2010), and all the double mutants seeds lacked the ray-like structures that were observed around the wild type (Fig. 2d). Among the single mutants, only *muci21* and *cesa5* displayed clear ray-like structures (Fig. 2d), while *sos5* only had more diffuse cellulose as previously shown (Fig. 2d; Griffiths et al., 2014). The impact of the different mutant combinations on cellulose architecture were also supported by crystalline polymer birefringence (Fig. S1c). In short, *CESA5, SOS5,* or *MUCI21* were epistatic to *MUCI10* for pectin adherence (Fig. 2b,c), via partially overlapping mechanisms, and the loss of any two players severely impaired cellulose structure. This double mutant analysis highlights the genetic complexity of cell wall biosynthesis in the seed coat and reveals how extracellular polysaccharide organization can be dramatically reshaped when more than one structural component is modified.

### The elongation and substitution of HM modulate salt tolerance

The newly generated mutant collection affecting multiple classes of wall polymers enabled us to investigate the physiological consequences of altering mucilage structure. We established a novel seed germination and salt stress assay using aqueous solutions in 24-well plates. Nearly all wild-type and mutant seeds imbibed in water germinated within 24 h post-stratification (Fig. 3a). However, when placed in 150 mM CaCl_2_, few wild-type seeds germinated even after 48 h of exposure to constant light. We initially hypothesized that mucilage-defective mutants might be more susceptible to salt stress, but unexpectedly found that *muci10* and *muci10 muci21* seeds had over 5-fold higher germination rate at this stage (Fig. 3a). The other mutant combinations germinated like the wild type at all time points. Only *muci10* and *muci10 muci21* had significantly longer radicles at 72 h in 150 mM CaCl_2_ (Fig. 3b and Fig. 3d), even though most mutants had around a two-fold higher flotation rate compared to the wild type (Fig. 3c). The enhanced germination rate and radicle growth of *muci10* in 150 mM CaCl_2_ was replicated in multiple assays, including up to 100 seeds per well and independent growth batches (Fig. 3e-g).

**Fig. 3.**
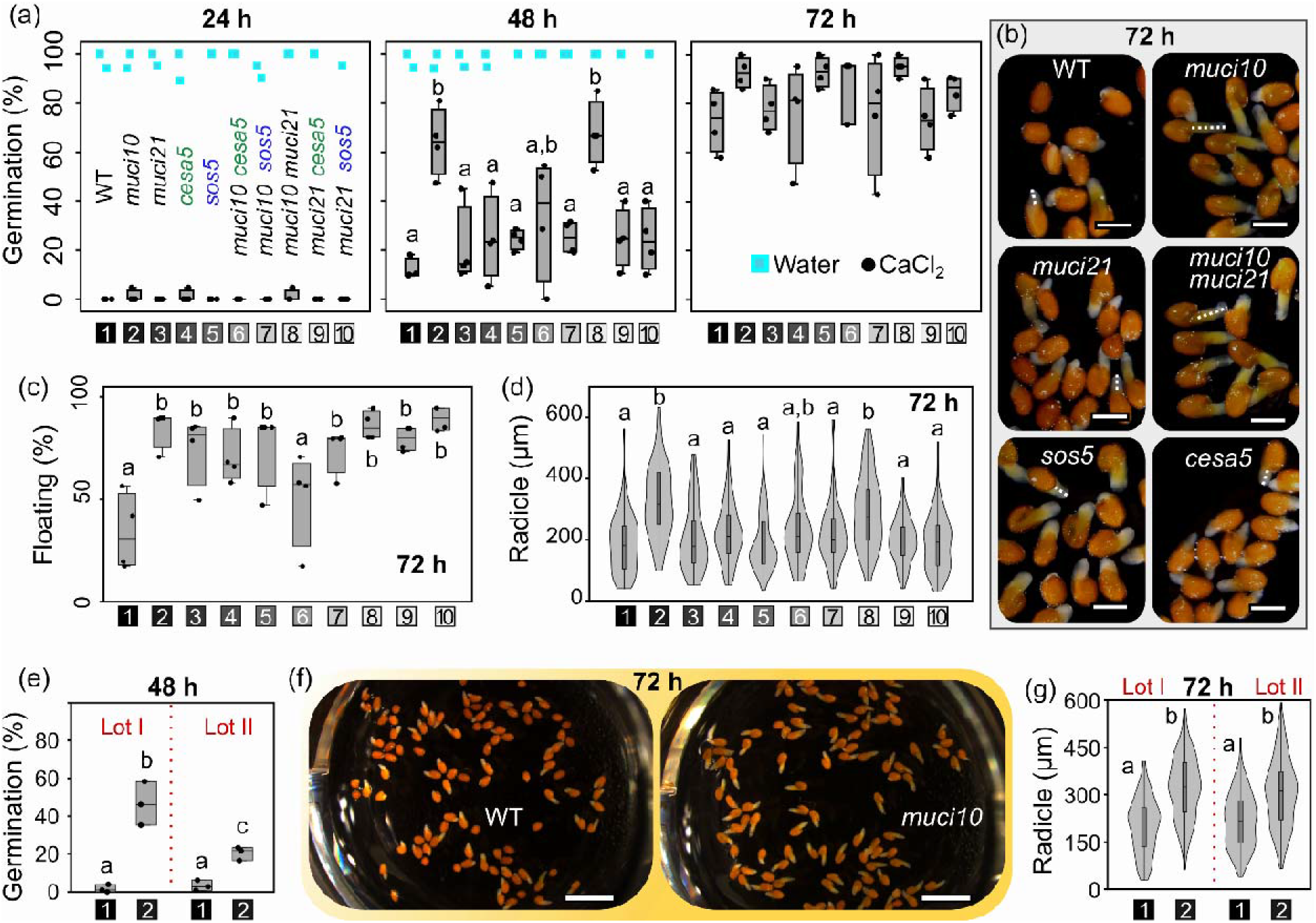
Germination of seeds in water and CaCl_2_. (a) Germination of stratified seeds. Box plots show germination of single and double mutants (4 plants, ~20 seeds each, per genotype) treated with 150 mM CaCl_2_. In water, nearly all seeds germinated within 24 h. (b) to (d) Further analyses of seeds from (a) in the CaCl_2_ treatment at 72 h. (b) Representative images of germinated seeds, with dashed lines indicating radicle length. (c) Box plots of seed flotation. (d) Violin and box plots of the radicle lengths. (e) to (g) Elevated *muci10* tolerance to 150 mM CaCl_2_ stress compared to the wild type (WT) was validated using larger quantities of seeds from two independent growth batches. (e) Germination rates at 48 h (3 plants, with ~100 seeds each) per genotype and seed lot. (f) Images of wells from the first seed lot at 72 h. (f) Radicle growth in CaCl_2_ in two seed lots. All X-axes are labeled using the legend in (a), and letters mark significant changes (one-way ANOVA with Tukey test, P < 0.05). Scale bars = 0.5 mm (b) and **2** mm (f).

To evaluate the basis of the observed salt tolerance, we assayed the effects of the *muci10* mutation in additional stress conditions. The use of 150 mM NaCl also reduced the rate of seed germination, but radicles that protruded from NaCl-treated seeds failed to further elongate compared to the CaCl_2_ treatment (Fig. S2). Nevertheless, *muci10* and *muci10 muci21* germinated faster than wild type in both salt treatments (Fig. 3a and Fig. S3a). All seeds sunk in water within the stratification period (Fig. S3b), but a significant proportion of certain seeds (only *muci21* in NaCl, and most mutants in CaCl_2_) continued to float in the salt solutions (Fig. S3c). When subjected to ionic (150 mM CaCl_2_ or MgCl_2_) or purely osmotic stress (PEG 4000 or sorbitol) of equivalent pressure, the germination rate of *muci10* seeds was significantly higher than wild type only in calcium salt stress (Fig. 4a). Once protruded from the seed coat, *muci10* radicles elongated significantly faster than wild type in both CaCl_2_ and sorbitol treatments (Fig. 4b; despite 3-fold difference in sample sizes), while the magnesium and PEG solutions showed higher toxicity to both genotypes. Overall, unbranched HM mutant seeds primarily tolerated high amounts of Ca^2+^ cations, which can cross-link unesterified pectin (Voiniciuc *et al.*, 2015c; Šola *et al.*, 2019). Switching water and 150 mM CaCl_2_ solutions at 24 h post-stratification demonstrated that *muci10* enhances growth in calcium stress during radicle emergence as well as subsequent elongation (Fig. S3d,e).

**Fig. 4.**
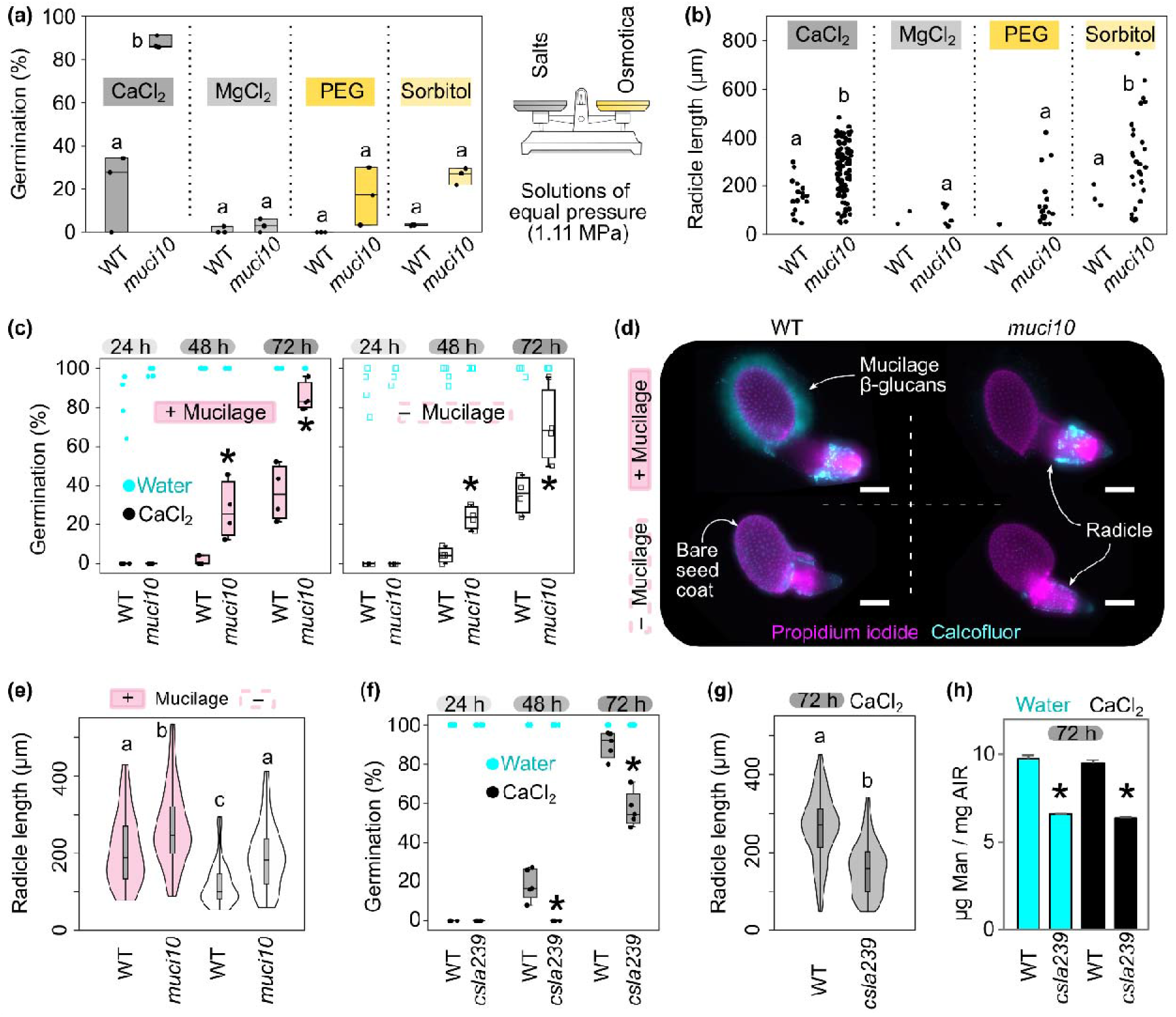
Dissecting how HM structure impacts germination in adverse conditions. (a) Germinati**on** rates at 72 h post-stratification in 150 mM CaCl_2_, MgCl_2_ or two osmotica (PEG 4000 and sorbitol), with an equal osmotic pressure. (b) Jitter plots showing radicle lengths at 72 h, from the seeds that germinated in (a). (c) Germination of seeds with (+) or without (–; mill-extracted) mucilage. (d) Dual cell wall staining of seeds germinated at 72 h in CaCl_2_. All *muci10* seeds as well as mill-extracted wild-type (WT) seeds lack mucilage β-glucans. Scale bars = 200 μm. (e) Radicle lengths of seeds from (c) at 72 h in CaCl_2_. (f) Germination of WT and *csla239* triple mutant. (g) Radicle length of *csla239* is reduced compared to WT. Data is shown from three biological replicates in (a) and (b), or four biological replicates in (c) to (g). (h) Mannose content in germinated seeds shown as mean + SD of two technical replicates. In all panels, significant changes are marked by different letters (one-way ANOVA with Tukey test, P < 0.05) or asteris**ks** (student’s t-test, P < 0.05; compared to the corresponding WT).

We then investigated how mucilage removal impacts salt tolerance, by extracting seed coat polysaccharides using a ball mill prior to stratification. With or without mucilage, CaCl_2_-treated *muci10* seeds germinated faster than wild-type (Fig. 4c). Mucilage β-glucans continue to encapsulate wild-type seeds at 72 h post-stratification (Fig. 4d), but were absent from de-mucilaged wild-type seeds and from HM-deficient *muci10* seeds (regardless of treatment). Despite not altering the germination rates of after-ripened seeds, the mucilage extraction significantly reduced the radicle length of each genotype compared to the intact controls (Fig. 4e). To evaluate the roles of different enzymes involved in HM biosynthesis, we then compared the germination rates of *muci10* and *csla2-3* (Fig. S4), which have similar mucilage defects (Voiniciuc *et al.*, 2015b). CaCl_2_-treated *csla2* resembled the wild type, but the mannose content of *csla2* germinated seeds was reduced by only 7% (t-test, P < 0.05) in either water or CaCl_2_ (Fig. S4c and Table S4), suggesting that additional CSLAs elongate HM in the same tissues. Using microarray data, we found that the transcription of *CSLA2, CSLA3, CSLA9* along with *CSLA7* and *CSLA11* (to a lesser extent) increased during germination relative to dry seeds (Fig. S4d). Compared to the wild type, we found that the *csla2-1 csla3-2 csla9-1* triple mutant (abbreviated as *csla239),* reported to have glucomannan-deficient stems (Goubet *et al.*, 2009), had significantly lower germination (Fig. 4f) and smaller radicles (Fig. 4g) in the CaCl_2_ treatment. The *csla239* triple mutant reduced the mannan content of germinated seeds by one-third (Fig. 4h and Table S5), indicating that even a partial reduction of HM elongation significantly impaired growth under salt stress. Since a *csla7* mutant was defective in embryogenesis (Goubet *et al.*, 2003, 2009), we expect that the full disruption of HM elongation in seeds would be lethal.

In summary, we found that the biosynthesis of two substituted hemicelluloses in the seed coat epidermis can be uncoupled and that HM and xylan have largely independent functions. HM substituted by MUCI10 is responsible for controlling pectin density, supporting cellulose synthesis and modulating seed tolerance to salt stress. In contrast, *MUCI21, CESA5* and *SOS5* are all epistatic to *MUCI10* for pectin adherence to the seed surface, via partially overlapping means (Fig. 1a). Since *muci21, cesa5* and *sos5* had additive effects (Fig. 1, Fig. 2, and Fig. S1, *cesa5 sos5* from Griffiths *et al.*, 2014, 2016), Arabidopsis seed mucilage structure must be controlled by a genetic network that is more complex than its carbohydrate composition suggests. For instance, the disruption of HM biosynthesis (Fig. 2; Yu *et al.*, 2014; Voiniciuc *et al.*, 2015b) or of cortical microtubule organization (Yang *et al.*, 2019) reduces the distribution of cellulose but not mucilage adherence. Our analysis of *muci10* and *cesa5* single and double mutants indicates that the cellulosic microfibrils essential for pectin attachment might be closer to the seed surface than previously thought (see remnants of rays in Fig. 2d and Fig. S1c).

In addition to gaining insight into the genetic regulation of mucilage properties, we discovered that HM structure modulates seed germination in CaCl_2_ solutions, and to a lesser extent in other ionic/osmotic conditions. Ca^2+^ ions can cross-link unesterified mucilage pectin and all the generated double mutants had elevated flotation compared to the wild type. However, only the *muci10* mutation promoted germination in CaCl_2_, while the *csla239* triple mutant reduced it. Consistent with these effects, *MUCI10* and other HM biosynthetic genes were up-regulated during seed germination (Fig. 1d and Fig. S4d), while *MUCI21* was not. Since *CESA5* was also expressed in germinating seeds (Fig. 1d) and *sos5* roots are overly sensitive to salt (no ATH1 microarray probe; Basu *et al.*, 2016), *muci10 cesa5* and *muci10 sos5* double mutants may offset the benefit of *muci10* (Fig. 3). The presence of unbranched HM could directly alter the ability of cell walls to expand under salt stress. HM deficiencies also modify pectin properties, by lowering the degree of methylesterification (Yu *et al.*, 2014), so *muci10* mucilage might be able to sequester calcium ions that would otherwise inhibit the expansion of inner cell layers.

In addition, unsubstituted HM in *muci10* should be more readily hydrolyzed or transglycosylated by β-1,4-mannanases (MAN; Schröder *et al.*, 2009), which are expressed during Arabidopsis seed imbibition (Fig. 1d,e). Mutations in *MAN5, MAN7,* and particularly *MAN6* are known to reduce germination in favorable conditions (Iglesias-Fernández *et al.*, 2011). We hypothesize that MAN enzymes might directly alter cell wall expansion, mobilize energy reserves and/or release an HM-derived molecular signal to enhance salt tolerance. Only water-treated seedlings accumulated large amounts of glucose (Tables S4 and S5), likely derived from starch. Seeds germinating in salt stress might need to mobilize carbon reserves from HM and potentially other mucilage polymers to sustain growth (Fig. 4). HM structure varies extensively in natural Arabidopsis populations (Voiniciuc *et al.*, 2016), so it might already modulate how seeds disperse, germinate and tolerate brackish waters containing hostile levels of Ca^2+^ and/or Na^+^. Consistent with this hypothesis, the constitutive expression of an enzyme involved in producing GDP-mannose, the sugar donor for HM elongation, elevated the mannose content of Arabidopsis seedlings and their tolerance to 150 mM NaCl (He *et al.*, 2017). Since the world faces rising sea levels and the expansion of saline environments, engineering salt tolerance remains a major challenge in crop production.

In conclusion, we have deciphered the contrasting roles of two classes of hemicelluloses in establishing seed mucilage properties and demonstrated new roles for HM elongation and substitution in radicle emergence as well as elongation during calcium salt stress. The multiwell cultivation system established in this study can be used to explore the physiological consequences of additional cell wall modifications. The overlapping expression profiles of multiple HM-related genes (Fig. 1d and S4d) highlights the need to investigate the specificity of these players on the cellular level in future studies. Future studies using *in vitro* (Liepman *et al.*, 2005; Yu *et al.*, 2018) or synthetic biology (Voiniciuc *et al.*, 2019) approaches are required to elucidate the glycan structures yielded by different enzyme isoforms, or combinations thereof.

## Accession Numbers

MUCI10 (At2g22900); MUCI21 (At3g10320); CESA5 (At5g09870); SOS5 (At3g46550); CSLA2 (At5g22740); CSLA3 (At1g23480); CSLA9 (At5g03760)

## Acknowledgements

We thank Stefanie Müller, Benita Schmitz and Stefanie Clauß for excellent technical assistance with plant cultivation. We are also grateful to Dr. Debora Gasperini and the Imaging Unit at the Leibniz Institute of Plant Biochemistry for microscope access. The *csla239* triple mutant was kindly provided by Professor Paul Dupree (University of Cambridge). The research was funded in part by the Natural Sciences and Engineering Research Council of Canada (NSERC PGS-D3 to C.V.), Deutsche Forschungsgemeinschaft (DFG research grant 414353267 to C.V.; and US98/13-1 to B.U.) and by the Ministry of Innovation, Science and Research of North-Rhine Westphalia within the framework of the NRW Strategieprojekt BioSC (No. 313/323 400 00213 to B.U.). Generation of the CCRC series of monoclonal antibodies used in this work was supported by a grant from the NSF Plant Genome Program (DBI-0421683).

## Author Contributions

B.Y. and C.V. designed the research. B.U. and C.V. supervised the first and second halves of the project, respectively. B.Y., F.H. and C.V. performed experiments and data analysis. C.V. wrote the article using drafts from B.Y. and valuable feedback from B.U.

## Supporting Information

Additional Supporting Information is found in the online version of this article

**Fig. S1** Xylan and crystalline cellulose distribution around seeds.

**Fig. S2** Morphology of seeds in CaCl_2_ and NaCl treatments.

**Fig. S3** Seed germination and flotation rates in water and salt stress.

**Fig. S4** Roles of heteromannan-related genes during seed germination.

**Table S1** Insertional mutants and genotyping primers used in this study.

**Table S2** Monosaccharide composition of total mucilage extracted from seeds.

**Table S3** Detachment of mucilage components after gentle shaking.

**Table S4** Cell wall composition of *csla2* mutant germinated seeds.

**Table S5** Cell wall composition of *muci10* and *csla239* germinated seeds.

## References

Anderson CT, Carroll A, Akhmetova L, Somerville C. 2010. Real-Time Imaging of Cellulose Reorientation during Cell Wall Expansion in Arabidopsis Roots. Plant Physiology 152: 787–796.

Basu D, Tian L, Debrosse T, Poirier E, Emch K, Herock H, Travers A, Showalter AM. 2016. Glycosylation of a Fasciclin-Like Arabinogalactan-Protein (SOS5) Mediates Root Growth and Seed Mucilage Adherence via a Cell Wall Receptor-Like Kinase (FEI1/FEI2) Pathway in Arabidopsis. PLOS ONE 11: e0145092.

Ben□Tov D, Idan□Molakandov A, Hugger A, Ben□Shlush I, Günl M, Yang B, Usadel B, Harpaz□Saad S. 2018. The role of COBRA LIKE 2 function, as part of the complex network of interacting pathways regulating Arabidopsis seed mucilage polysaccharide matrix organization. The Plant Journal 94: 497–512.

Berendzen K, Searle I, Ravenscroft D, Koncz C, Batschauer A, Coupland G, Somssich IE, Ülker B. 2005. A rapid and versatile combined DNA/RNA extraction protocol and its application to the analysis of a novel DNA marker set polymorphic between Arabidopsis thaliana ecotypes Col-0 and Landsberg erecta. Plant Methods 1: 4.

Goubet F, Barton CJ, Mortimer JC, Yu X, Zhang Z, Miles GP, Richens J, Liepman AH, Seffen K, Dupree P. 2009. Cell wall glucomannan in Arabidopsis is synthesised by CSLA glycosyltransferases, and influences the progression of embryogenesis. The Plant Journal 60: 527–538.

Goubet F, Misrahi A, Park SK, Zhang Z, Twell D, Dupree P. 2003. AtCSLA7, a Cellulose Synthase-Like Putative Glycosyltransferase, Is Important for Pollen Tube Growth and Embryogenesis in Arabidopsis. Plant Physiology 131: 547–557.

Griffiths JS, Crepeau M-J, Ralet M-C, Seifert GJ, North HM. 2016. Dissecting Seed Mucilage Adherence Mediated by FEI2 and SOS5. Frontiers in Plant Science 7: 1–13.

Griffiths JS, North HM. 2017. Sticking to cellulose: exploiting Arabidopsis seed coat mucilage to understand cellulose biosynthesis and cell wall polysaccharide interactions. New Phytologist 214: 959–966.

Griffiths JS, Šola K, Kushwaha R, Lam P, Tateno M, Young R, Voiniciuc C, Dean G, Mansfield SD, DeBolt S, et al. 2015. Unidirectional Movement of Cellulose Synthase Complexes in Arabidopsis Seed Coat Epidermal Cells Deposit Cellulose Involved in Mucilage Extrusion, Adherence, and Ray Formation. Plant Physiology 168: 502–520.

Griffiths JS, Tsai AYL, Xue H, Voiniciuc C, Šola K, Seifert GJ, Mansfield SD, Haughn GW. 2014. SALT-OVERLY SENSITIVE5 mediates arabidopsis seed coat mucilage adherence and organization through pectins. Plant Physiology 165: 991–1004.

Hammer Ø, Harper DAT, Ryan PD. 2001. PAST: Paleontological statistics software package for education and data analysis. Palaeontologia Electronica 4.

Harpaz-Saad S, McFarlane HE, Xu S, Divi UK, Forward B, Western TL, Kieber JJ. 2011. Cellulose synthesis via the FEI2 RLK/SOS5 pathway and CELLULOSE SYNTHASE 5 is required for the structure of seed coat mucilage in Arabidopsis. The Plant Journal 68: 941–953.

He C, Yu Z, Teixeira Da Silva JA, Zhang J, Liu X, Wang X, Zhang X, Zeng S, Wu K, Tan J, et al. 2017. DoGMP1 from Dendrobium officinale contributes to mannose content of water-soluble polysaccharides and plays a role in salt stress response. Scientific Reports 7: 1–13.

Iglesias-Fernández R, Rodríguez-Gacio MC, Barrero-Sicilia C, Carbonero P, Matilla A. 2011. Three endo-β-mannanase genes expressed in the micropylar endosperm and in the radicle influence germination of Arabidopsis thaliana seeds. Planta 233: 25–36.

Liepman AH, Wilkerson CG, Keegstra K. 2005. Expression of cellulose synthase-like (Csl) genes in insect cells reveals that CslA family members encode mannan synthases. Proceedings of the National Academy of Sciences of the United States of America 102: 2221–2226.

Mendu V, Griffiths JS, Persson S, Stork J, Downie AB, Voiniciuc C, Haughn GW, DeBolt S. 2011. Subfunctionalization of Cellulose Synthases in Seed Coat Epidermal Cells Mediates Secondary Radial Wall Synthesis and Mucilage Attachment. Plant Physiology 157: 441–453.

Money NP. 1989. Osmotic Pressure of Aqueous Polyethylene Glycols. Plant Physiology.

Narsai R, Law SR, Carrie C, Xu L, Whelan J. 2011. In-Depth Temporal Transcriptome Profiling Reveals a Crucial Developmental Switch with Roles for RNA Processing and Organelle Metabolism That Are Essential for Germination in Arabidopsis. Plant Physiology 157: 1342–1362.

North HM, Berger A, Saez-Aguayo S, Ralet M-C. 2014. Understanding polysaccharide production and properties using seed coat mutants: future perspectives for the exploitation of natural variants. Annals of Botany 114: 1251–1263.

Ralet M-C, Crépeau M-J, Vigouroux J, Tran J, Berger A, Sallé C, Granier F, Botran L, North HM. 2016. Xylans Provide the Structural Driving Force for Mucilage Adhesion to the Arabidopsis Seed Coat. Plant Physiology 171: 165–178.

Ruprecht C, Bartetzko MP, Senf D, Dallabernadina P, Boos I, Andersen MCF, Kotake T, Knox JP, Hahn MG, Clausen MH, et al. 2017. A synthetic glycan microarray enables epitope mapping of plant cell wall glycan-directed antibodies. Plant Physiology 175: 1094–1104.

Saez-Aguayo S, Rondeau-Mouro C, Macquet A, Kronholm I, Ralet MC, Berger A, Sallé C, Poulain D, Granier F, Botran L, et al. 2014. Local Evolution of Seed Flotation in Arabidopsis. PLoS Genetics 10: 13–15.

Schindelin J, Arganda-Carreras I, Frise E, Kaynig V, Longair M, Pietzsch T, Preibisch S, Rueden C, Saalfeld S, Schmid B, et al. 2012. Fiji: an open-source platform for biological-image analysis. Nature Methods 9: 676–682.

Schröder R, Atkinson RG, Redgwell RJ. 2009. Re-interpreting the role of endo-β-mannanases as mannan endotransglycosylase/hydrolases in the plant cell wall. Annals of Botany 104: 197–204.

Šola K, Dean GH, Haughn GW. 2019. Arabidopsis Seed Mucilage: A Specialised Extracellular Matrix that Demonstrates the Structure–Function Versatility of Cell Wall Polysaccharides. Annual Plant Reviews online 2: 1085–1116.

Sullivan S, Ralet M-C, Berger A, Diatloff E, Bischoff V, Gonneau M, Marion-Poll A, North HM. 2011. CESA5 Is Required for the Synthesis of Cellulose with a Role in Structuring the Adherent Mucilage of Arabidopsis Seeds. Plant Physiology 156: 1725–1739.

Voiniciuc C. 2016. Quantification of the Mucilage Detachment from Arabidopsis Seeds. BIO-PROTOCOL 6: 1–9.

Voiniciuc C. 2017. Whole-seed Immunolabeling of Arabidopsis Mucilage Polysaccharides. BIO-PROTOCOL 7: 1–10.

Voiniciuc C, Dama M, Gawenda N, Stritt F, Pauly M. 2019. Mechanistic insights from plant heteromannan synthesis in yeast. Proceedings of the National Academy of Sciences 116: 522–527.

Voiniciuc C, Engle KA, Günl M, Dieluweit S, Schmidt MH-W, Yang J-Y, Moremen KW, Mohnen D, Usadel B. 2018. Identification of Key Enzymes for Pectin Synthesis in Seed Mucilage. Plant Physiology 178: 1045–1064.

Voiniciuc C, Günl M. 2016. Analysis of Monosaccharides in Total Mucilage Extractable from Arabidopsis Seeds. BIO-PROTOCOL 6: 1–12.

Voiniciuc C, Günl M, Schmidt MH-W, Usadel B. 2015a. Highly Branched Xylan Made by IRREGULAR XYLEM14 and MUCILAGE-RELATED21 Links Mucilage to Arabidopsis Seeds. Plant physiology 169: 2481–95.

Voiniciuc C, Schmidt MH-W, Berger A, Yang B, Ebert B, Scheller H V., North HM, Usadel B, Günl M. 2015b. MUCILAGE-RELATED10 Produces Galactoglucomannan That Maintains Pectin and Cellulose Architecture in Arabidopsis Seed Mucilage. Plant Physiology 169: 403–420.

Voiniciuc C, Yang B, Schmidt MH-W, Günl M, Usadel B. 2015c. Starting to Gel: How Arabidopsis Seed Coat Epidermal Cells Produce Specialized Secondary Cell Walls. International Journal of Molecular Sciences 16: 3452–3473.

Voiniciuc C, Zimmermann E, Schmidt MH-W, Günl M, Fu L, North HM, Usadel B. 2016. Extensive Natural Variation in Arabidopsis Seed Mucilage Structure. Frontiers in Plant Science 7: 1–14.

Western TL. 2012. The sticky tale of seed coat mucilages: production, genetics, and role in seed germination and dispersal. Seed Science Research 22: 1–25.

Yang B, Voiniciuc C, Fu L, Dieluweit S, Klose H, Usadel B. 2019. TRM4 is essential for cellulose deposition in Arabidopsis seed mucilage by maintaining cortical microtubule organization and interacting with CESA3. New Phytologist 221: 881–895.

Yu L, Lyczakowski JJ, Pereira CS, Kotake T, Yu X, Li A, Mogelsvang S, Skaf MS, Dupree P. 2018. The Patterned Structure of Galactoglucomannan Suggests It May Bind to Cellulose in Seed Mucilage. Plant Physiology 178: 1011–1026.

Yu L, Shi D, Li J, Kong Y, Yu Y, Chai G, Hu R, Wang J, Hahn MG, Zhou G. 2014. CELLULOSE SYNTHASE-LIKE A2, a glucomannan synthase, is involved in maintaining adherent mucilage structure in arabidopsis seed. Plant Physiology 164: 1842–1856.

Zhong R, Cui D, Phillips DR, Ye Z-H. 2018. A Novel Rice Xylosyltransferase Catalyzes the Addition of 2-O-Xylosyl Side Chains onto the Xylan Backbone. Plant and Cell Physiology 59: 554–565.

