## Supplemental Information for "Seed hemicelluloses tailor mucilage properties and salt tolerance"

The following Supporting Information is available for this article:

**Fig. S1** Xylan and crystalline cellulose distribution around seeds.

**Fig. S2** Morphology of seeds in $CaCl_2$ and NaCl treatments.

**Fig. S3** Seed germination and flotation rates in water and salt stress.

**Fig. S4** Roles of heteromannan-related genes during seed germination.

**Table S1** Insertional mutants and genotyping primers used in this study.

**Table S2** Monosaccharide composition of total mucilage extracted from seeds.

**Table S3** Detachment of mucilage components after gentle shaking.

**Table S4** Cell wall composition of *csla2* mutant germinated seeds.

**Table S5** Cell wall composition of *muci10* and *csla239* germinated seeds.

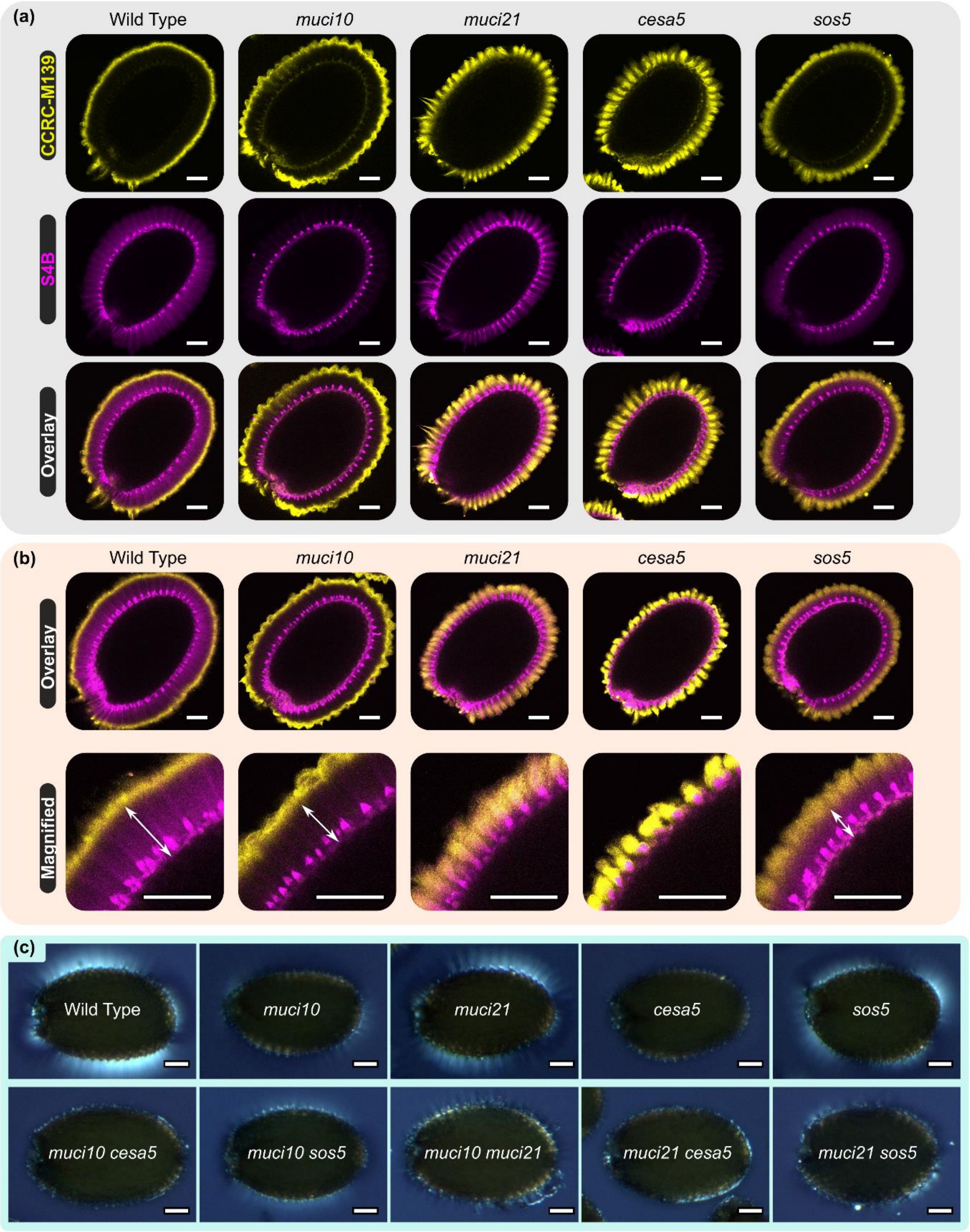


**Figure S1**. Xylan and crystalline cellulose distribution around seeds.
(a) Immunolabeling of seeds with the CCRC-M139 antibody, which binds to stretches of unbranched xylan (Ruprecht et al., 2017). Cellulose was counter-stained with S4B. (b) Overlay of CCRC-M139 and S4B signals for another seed per genotype. In the magnified view, arrows indicate the distance between the xylan epitopes and the seed surface. (c) Birefringence of crystalline structures around the seed surface, viewed under polarized light. Bars = 100 µm.

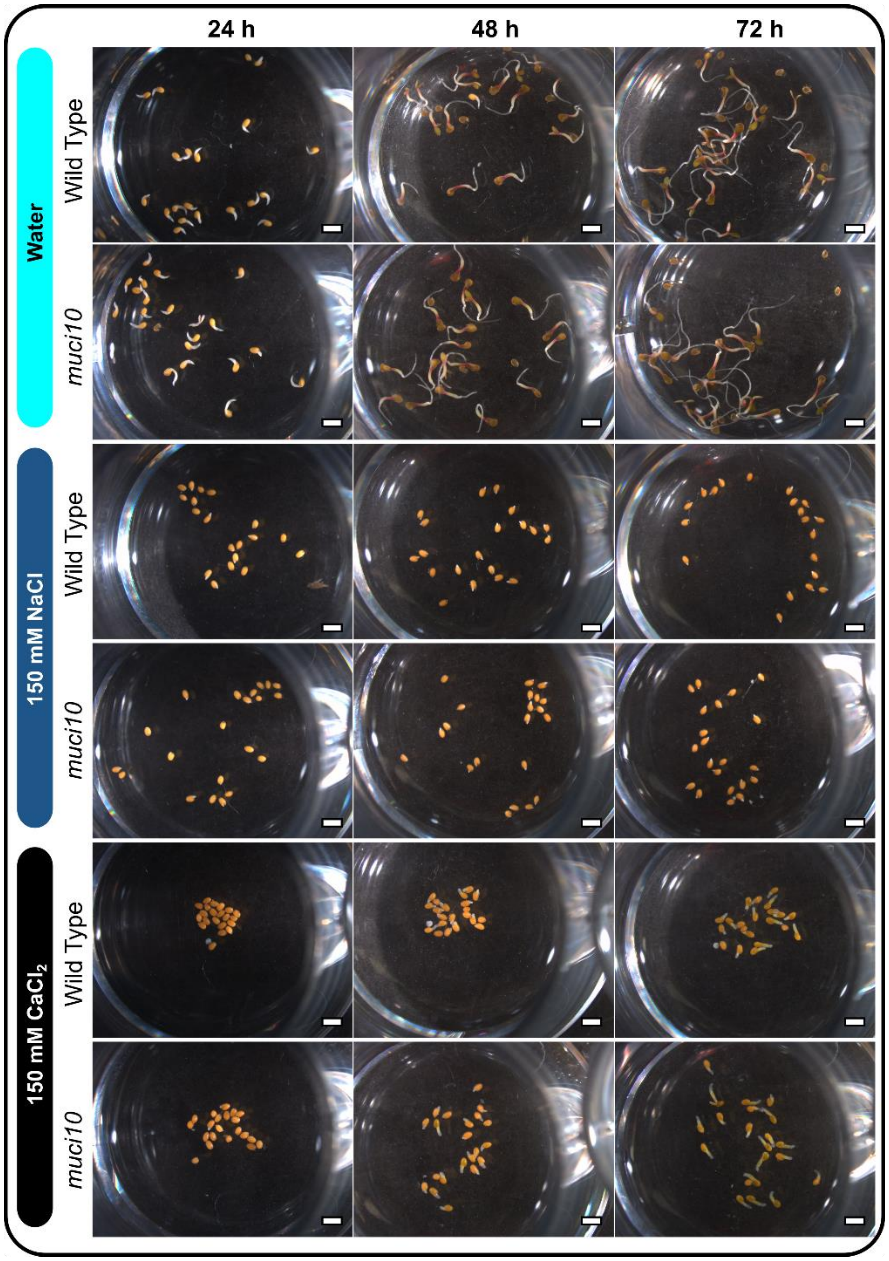


**Figure S2**. Morphology of seeds in $CaCl_2$ and NaCl treatments.
Images of seeds at three timepoints post-stratification in water, 150 mM NaCl, or 150 mM $CaCl_2$. Scale bars = 1 mm.

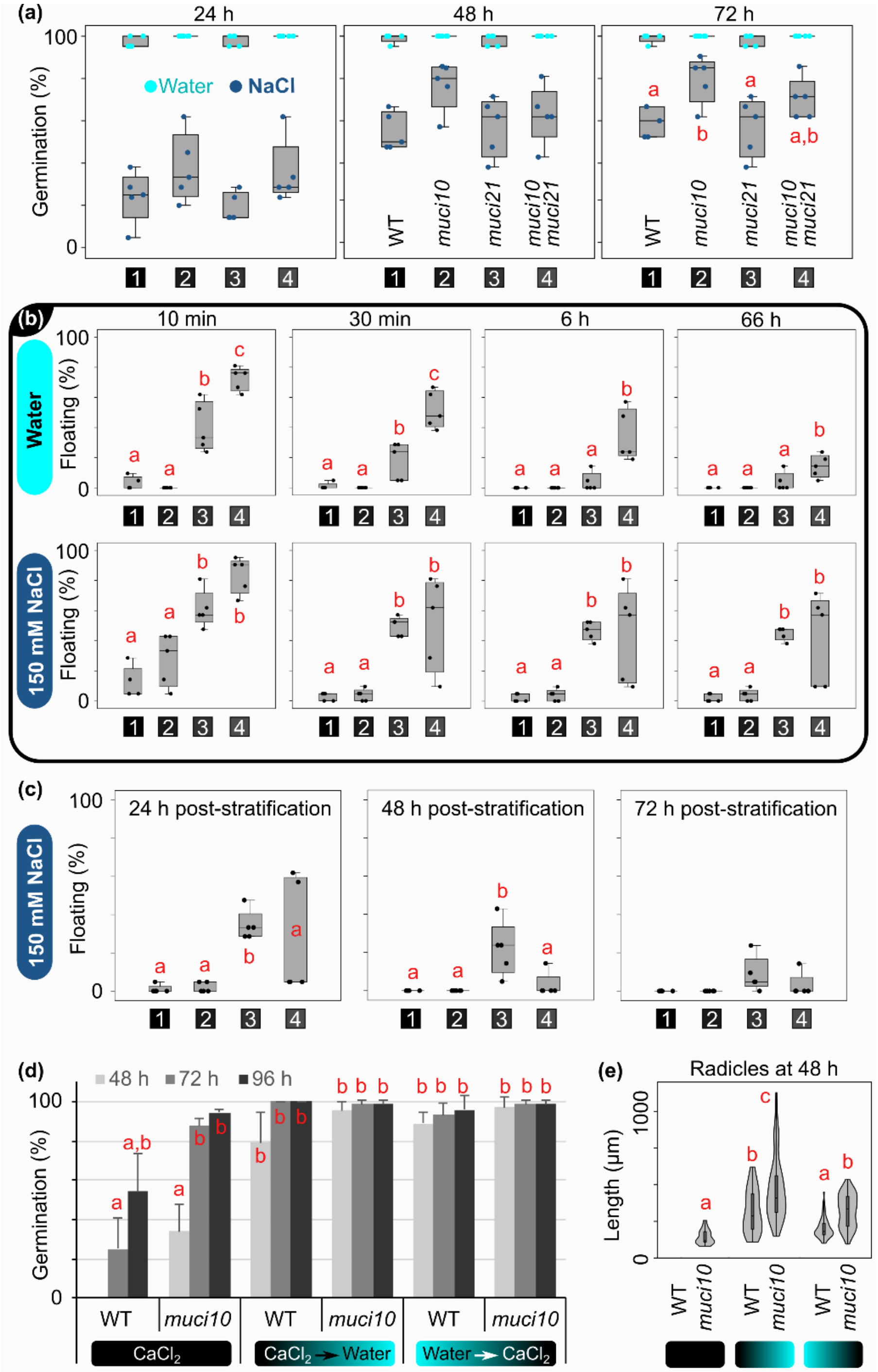


**Figure S3**. Seed germination and flotation rates in water and salt stress.
(a) Germination of seeds at three time points, after stratification in water or 150 mM NaCl. (b) Seed flotation in water or NaCl during the stratification period. (c) Flotation of stratified seeds after transfer to constant light. All box plots show five biological replicates. Letters mark significant changes (one-way ANOVA with Tukey test, $P < 0.05$). (d) Germination rates (mean + SD of three biological replicates) in 150 mM $CaCl_2$ or following solution exchange at 24 h post-stratification. (e) Violin and box plots of radicle length from seeds that germinated in (d) at 48 h post-stratification. The treatments are colored according to the legend in (d).

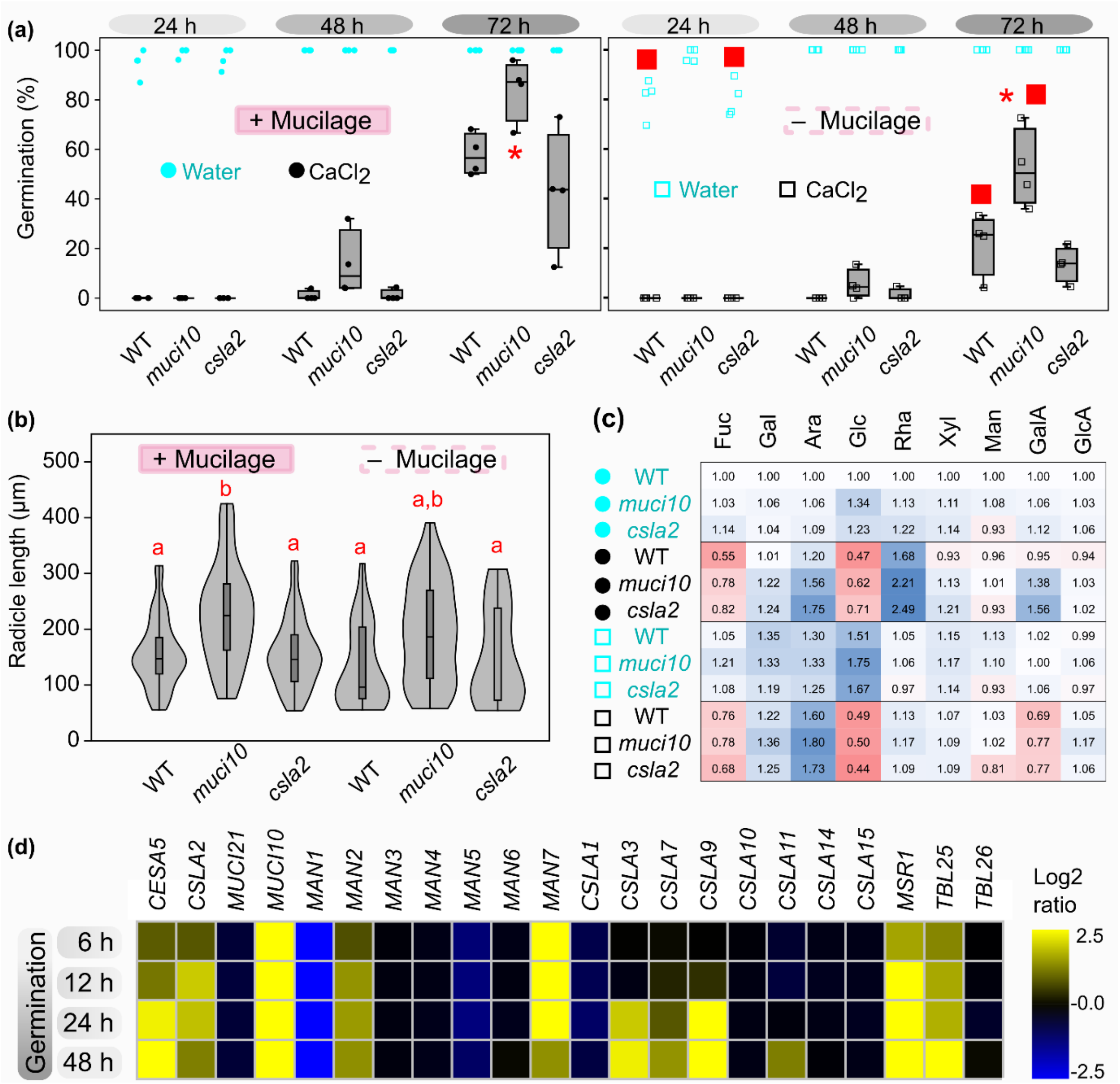


**Figure S4**. Roles of heteromannan-related genes during seed germination.
(a) Post-stratification, germination rates of intact seeds (+ Mucilage) or after total mucilage extraction (- Mucilage). (b) Violin and box plots showing radicle length of seeds (intact or de-mucilaged) at 72 h post-stratification in 150 mM $CaCl_2$. (c) Heatmap of absolute monosaccharide abundance of germinated seeds at 72 h post-stratification, showing fold changes relative to the wild type seeds treated with water. Four biological replicates per genotype were examined in (a) to (c), at two weeks post-harvest. Significant changes are marked by different letters (one-way ANOVA with Tukey test, $P < 0.05$) or asterisks (student's *t*-test, $P < 0.05$; compared to the corresponding WT). Red squares mark significant effect of mucilage extraction relative to the intact seed control (student's *t*-test, $P < 0.05$). (d) GENEVESTIGATO heatmap of transcriptional changes (log2 ratio) in germinating seeds post-stratification, relative to dry seeds. Heteromannan-related genes are involved in polysaccharide elongation (CSLAs, MSR), galactosylation (MUCI10), hydrolysis/transglycosylation (MANs), and acetylation (TBLs).

**Table S1**. Insertional mutants and genotyping primers used in this study.
Single mutants analyzed in this study, and the 5' to 3' sequences of the gene-specific primers used together with the SALK T-DNA primer LBb1.3 (ATTTTGCCGATTTCGGAAC). All primers were designed using http://signal.salk.edu/tdnaprimers.2.html.

| Allele | T-DNA Line | Forward Primer | Reverse Primer |
|---|---|---|---|
| *muci10-1* | SALK_061576C | AAACCTACCAATCGAAACACG | TAAACCAAACCAGACAATGCC |
| *muci21-1* | SALK_041744 | AATGCGAAGTGATCCATGAAG | TGGCTCTTGTTTAAGCCATTC |
| *cesa5-1* | SALK_118491 | TCTCAAGCCATTTCAAACCAC | TTCATGATGGATCCTCAGTCG |
| *sos5-2* | SALK_125874 | GAAACTGGGAATAACCTTCGG | AGCTTCTCGAGACCAAACCTC |
| *csla2-3* | SALK_149092 | TAGATGGTCTTGTGGACCTGC | CAAAAGAACCCTTGGAGCTTC |
| *csla2-1* *csla3-2* *csla9-1* | SALK_065083 | ACCAGTTTACAAACCCGAACC | GGTGGAAGTGGAGTGTCAAAG |
| | SALK_087535 | AGGACACAAACCAAGATCTCG | CAGGATCAGGCTGAAAATCAG |
| | SALK_071916 | CATTTTCACTAGATCCGCCAC | TCCGGTACAAGAACTGTAGCG |

**Table S2**. Monosaccharide composition of total mucilage extracted from seeds.
Total mucilage was extracted from seeds using water and a ball mill. Data show the mean molar percentage ± SD of four biological replicates (only three for the *muci21 sos5* double mutant).

| | **Rhamnose** | **Arabinose** | **Galactose** | **Glucose** | **Xylose** | **Mannose** | **Galacturonic Acid** |
|---|---|---|---|---|---|---|---|
| Wild Type | 39.75 ± 3.22 | 1.63 ± 0.20 | 1.27 ± 0.16 | 0.78 ± 0.10 | 3.71 ± 0.35 | 0.85 ± 0.10 | 52.00 ± 4.04 |
| *muci10* | 43.49 ± 1.27 | 1.76 ± 0.08 | 0.68 ± 0.06 | 0.79 ± 0.14 | 4.42 ± 0.06 | 0.54 ± 0.10 | 48.32 ± 1.43 |
| *muci21* | 43.37 ± 1.90 | 1.50 ± 0.15 | 1.24 ± 0.08 | 0.89 ± 0.14 | 1.62 ± 0.07 | 0.91 ± 0.05 | 50.48 ± 2.04 |
| *cesa5* | 41.76 ± 1.77 | 1.50 ± 0.17 | 1.01 ± 0.09 | 0.98 ± 0.15 | 3.36 ± 0.28 | 0.79 ± 0.02 | 50.60 ± 2.14 |
| *sos5* | 41.43 ± 1.73 | 1.69 ± 0.28 | 1.36 ± 0.08 | 1.18 ± 0.39 | 3.64 ± 0.33 | 0.99 ± 0.09 | 49.71 ± 2.52 |
| *muci10 cesa5* | 41.83 ± 3.12 | 1.99 ± 0.14 | 0.95 ± 0.05 | 0.80 ± 0.32 | 3.62 ± 0.27 | 0.54 ± 0.05 | 50.27 ± 3.47 |
| *muci10 sos5* | 41.70 ± 2.69 | 1.93 ± 0.15 | 0.74 ± 0.03 | 0.61 ± 0.04 | 3.92 ± 0.50 | 0.53 ± 0.09 | 50.58 ± 3.32 |
| *muci10 muci21* | 42.14 ± 2.48 | 1.60 ± 0.12 | 0.60 ± 0.03 | 0.54 ± 0.05 | 1.62 ± 0.11 | 0.52 ± 0.04 | 52.99 ± 2.66 |
| *muci21 cesa5* | 42.29 ± 1.70 | 2.72 ± 0.41 | 1.55 ± 0.18 | 0.74 ± 0.12 | 2.19 ± 0.13 | 0.69 ± 0.08 | 49.82 ± 2.20 |
| *muci21 sos5* | 44.50 ± 1.20 | 1.89 ± 0.06 | 1.49 ± 0.02 | 1.04 ± 0.13 | 1.98 ± 0.11 | 1.03 ± 0.05 | 48.08 ± 1.45 |

**Table S3**. Detachment of mucilage components after gentle shaking.
Percentage of the total extractable mucilage monosaccharides that are non-adherent. Data show the mean ± SD of four biological replicates (only two for *sos5-2*).

| | **Rhamnose** | **Arabinose** | **Galactose** | **Glucose** | **Xylose** | **Mannose** | **Galacturonic Acid** |
|---|---|---|---|---|---|---|---|
| Wild Type | 75 ± 3 | 50 ± 5 | 19 ± 2 | 5 ± 2 | 68 ± 3 | 42 ± 1 | 71 ± 3 |
| *muci10* | 71 ± 4 | 46 ± 7 | 13 ± 3 | 5 ± 2 | 68 ± 4 | 51 ± 4 | 67 ± 4 |
| *muci21* | 92 ± 0 | 58 ± 7 | 21 ± 4 | 8 ± 6 | 51 ± 4 | 41 ± 4 | 90 ± 1 |
| *cesa5* | 89 ± 2 | 54 ± 4 | 27 ± 2 | 9 ± 2 | 86 ± 1 | 55 ± 5 | 86 ± 2 |
| *sos5* | 89 ± 1 | 39 ± 11 | 13 ± 5 | 9 ± 3 | 86 ± 2 | 48 ± 4 | 87 ± 0 |
| *muci10 cesa5* | 86 ± 4 | 38 ± 6 | 13 ± 3 | 5 ± 2 | 83 ± 3 | 46 ± 6 | 84 ± 6 |
| *muci10 sos5* | 87 ± 0 | 52 ± 6 | 16 ± 2 | 5 ± 1 | 85 ± 3 | 58 ± 3 | 84 ± 1 |
| *muci10 muci21* | 89 ± 2 | 47 ± 3 | 14 ± 1 | 6 ± 3 | 45 ± 1 | 44 ± 2 | 86 ± 1 |
| *muci21 cesa5* | 92 ± 1 | 32 ± 3 | 12 ± 2 | 4 ± 0 | 71 ± 1 | 35 ± 3 | 89 ± 3 |
| *muci21 sos5* | 94 ± 1 | 53 ± 5 | 23 ± 2 | 7 ± 1 | 75 ± 1 | 46 ± 2 | 90 ± 2 |

**Table S4**. Cell wall composition of *csla2* mutant germinated seeds.
Abundance of monosaccharides (µg/mg AIR) at 72 h post-stratification. Dry seeds were imbibed in water (DW) or $CaCl_2$ (DC). Alternatively, seeds were first de-mucilaged using a Retsch ball mill before water (RW) or $CaCl_2$ (RC) treatment. Data show the mean ± SD of two technical replicates from a pool of four biological replicates. GalA, galacturonic acid; GlcA, glucuronic acid.

| | | Fucose | Galactose | Arabinose | Glucose | Rhamnose | Xylose | Mannose | GalA | GlcA |
|---|---|---|---|---|---|---|---|---|---|---|
| DW | WT | 3.5 ± 0.0 | 44.8 ± 0.2 | 34.3 ± 0.2 | 82.8 ± 0.5 | 26.2 ± 0.3 | 16.4 ± 0.3 | 8.8 ± 0.0 | 40.1 ± 0.7 | 6.9 ± 0.9 |
| | *muci10* | 3.6 ± 0.1 | 47.5 ± 0.6 | 36.4 ± 1.1 | 111.1 ± 15.9 | 29.5 ± 0.5 | 18.3 ± 0.7 | 9.5 ± 0.1 | 42.4 ± 4.5 | 7.2 ± 0.1 |
| | *csla2* | 3.9 ± 0.6 | 46.7 ± 1.4 | 37.3 ± 2.7 | 102.1 ± 0.8 | 31.8 ± 1.4 | 18.8 ± 1.5 | 8.2 ± 0.0 | 45.1 ± 4.1 | 7.3 ± 0.5 |
| DC | WT | 1.9 ± 0.0 | 45.4 ± 2.9 | 41.0 ± 2.4 | 39.1 ± 2.1 | 43.8 ± 0.6 | 15.4 ± 0.7 | 8.5 ± 0.1 | 38.1 ± 5.5 | 6.5 ± 0.2 |
| | *muci10* | 2.7 ± 0.2 | 54.5 ± 8.4 | 53.5 ± 7.4 | 51.3 ± 5.2 | 57.9 ± 5.1 | 18.5 ± 1.7 | 8.9 ± 0.3 | 55.4 ± 10.1 | 7.2 ± 0.1 |
| | *csla2* | 2.8 ± 0.6 | 55.6 ± 13.0 | 60.2 ± 16.1 | 59.2 ± 17.3 | 65.2 ± 12.7 | 19.8 ± 2.8 | 8.2 ± 1.0 | 62.5 ± 2.5 | 7.1 ± 0.8 |
| RW | WT | 3.6 ± 0.4 | 60.5 ± 9.4 | 44.6 ± 6.9 | 125.0 ± 24.0 | 27.5 ± 1.7 | 18.9 ± 1.1 | 9.9 ± 0.4 | 40.9 ± 6.0 | 6.9 ± 0.1 |
| | *muci10* | 4.2 ± 0.5 | 59.7 ± 8.4 | 45.6 ± 6.4 | 145.2 ± 23.0 | 27.7 ± 1.6 | 19.2 ± 0.9 | 9.7 ± 0.3 | 40.1 ± 6.4 | 7.3 ± 0.9 |
| | *csla2* | 3.7 ± 0.4 | 53.5 ± 7.7 | 43.0 ± 5.6 | 138.6 ± 26.0 | 25.5 ± 1.5 | 18.7 ± 1.5 | 8.2 ± 0.2 | 42.4 ± 2.4 | 6.7 ± 0.5 |
| RC | WT | 2.6 ± 0.1 | 54.7 ± 3.7 | 54.7 ± 0.5 | 40.2 ± 3.8 | 29.5 ± 2.6 | 17.6 ± 0.0 | 9.0 ± 0.5 | 27.9 ± 0.8 | 7.3 ± 0.2 |
| | *muci10* | 2.7 ± 0.2 | 61.2 ± 6.2 | 61.9 ± 5.4 | 41.7 ± 2.7 | 30.7 ± 2.0 | 18.0 ± 1.0 | 9.0 ± 0.6 | 30.8 ± 2.9 | 8.1 ± 0.2 |
| | *csla2* | 2.4 ± 0.3 | 55.9 ± 2.5 | 59.4 ± 0.5 | 36.7 ± 0.1 | 28.4 ± 2.4 | 18.0 ± 0.3 | 7.1 ± 0.2 | 31.0 ± 0.8 | 7.3 ± 0.4 |

**Table S5**. Cell wall composition of *muci10* and *csla239* germinated seeds.
Abundance of monosaccharides (µg/mg AIR) at 72 h post-stratification. Dry seeds were imbibed in water (DW) or $CaCl_2$ (DC). Alternatively, seeds were first de-mucilaged using a Retsch ball mill before water (RW) or $CaCl_2$ (RC) treatment. Data show the mean ± SD of two technical replicates from a pool of four biological replicates. GalA, galacturonic acid; GlcA, glucuronic acid. Note that *muci10*, *csla239* and the corresponding wild-type (WT) seeds originate from two different plant growth batches, analyzed in parallel.

| | | **Fucose** | **Galactose** | **Arabinose** | **Glucose** | **Rhamnose** | **Xylose** | **Mannose** | **GalA** | **GlcA** |
|---|---|---|---|---|---|---|---|---|---|---|
| DW | WT | 5.0 ± 0.4 | 69.1 ± 6.3 | 53.3 ± 2.2 | 135.0 ± 10.4 | 38.7 ± 1.5 | 21.3 ± 0.5 | 10.3 ± 0.3 | 54.9 ± 3.2 | 8.2 ± 0.2 |
| | *muci10* | 5.1 ± 0.1 | 62.7 ± 3.0 | 49.2 ± 0.9 | 132.0 ± 6.0 | 33.5 ± 0.3 | 21.1 ± 0.2 | 9.5 ± 0.2 | 49.6 ± 1.0 | 7.7 ± 0.3 |
| DC | WT | 2.4 ± 0.1 | 57.1 ± 3.1 | 53.7 ± 0.8 | 36.9 ± 0.3 | 49.5 ± 0.7 | 17.2 ± 0.1 | 8.7 ± 0.3 | 48.5 ± 0.3 | 7.3 ± 0.3 |
| | *muci10* | 2.7 ± 0.2 | 56.0 ± 1.7 | 56.2 ± 1.8 | 45.2 ± 1.5 | 53.5 ± 1.6 | 18.5 ± 0.1 | 8.5 ± 0.2 | 56.4 ± 1.9 | 7.9 ± 0.1 |
| RW | WT | 4.8 ± 0.0 | 73.3 ± 3.9 | 57.8 ± 1.8 | 173.2 ± 6.0 | 31.2 ± 0.9 | 21.6 ± 0.8 | 10.1 ± 0.2 | 51.1 ± 2.9 | 9.0 ± 0.1 |
| | *muci10* | 5.1 ± 0.3 | 71.3 ± 4.4 | 55.8 ± 2.3 | 221.4 ± 12.6 | 31.2 ± 0.8 | 21.6 ± 0.4 | 10.1 ± 0.2 | 48.5 ± 3.4 | 8.6 ± 0.2 |
| RC | WT | 2.7 ± 0.2 | 64.8 ± 0.9 | 63.0 ± 3.9 | 41.9 ± 6.1 | 29.2 ± 3.4 | 18.5 ± 1.2 | 9.2 ± 0.4 | 31.2 ± 2.5 | 7.4 ± 0.1 |
| | *muci10* | 2.5 ± 0.1 | 61.7 ± 2.8 | 59.1 ± 1.0 | 33.9 ± 0.8 | 27.6 ± 0.0 | 17.4 ± 0.3 | 8.6 ± 0.1 | 30.3 ± 0.3 | 8.1 ± 0.2 |

| | | **Fucose** | **Galactose** | **Arabinose** | **Glucose** | **Rhamnose** | **Xylose** | **Mannose** | **GalA** | **GlcA** |
|---|---|---|---|---|---|---|---|---|---|---|
| DW | WT | 4.1 ± 0.0 | 57.0 ± 5.8 | 42.4 ± 1.6 | 172.7 ± 25.1 | 32.7 ± 1.0 | 18.9 ± 0.3 | 9.7 ± 0.2 | 46.4 ± 4.3 | 6.8 ± 0.2 |
| | *csla239* | 4.2 ± 0.2 | 57.6 ± 6.6 | 43.8 ± 3.1 | 145.8 ± 14.7 | 36.7 ± 2.4 | 19.5 ± 0.8 | 6.6 ± 0.0 | 54.9 ± 4.8 | 7.5 ± 0.6 |
| DC | WT | 2.8 ± 0.2 | 58.4 ± 2.4 | 54.7 ± 1.2 | 43.1 ± 0.3 | 51.0 ± 1.9 | 18.3 ± 0.2 | 9.5 ± 0.2 | 49.0 ± 2.5 | 7.9 ± 0.9 |
| | *csla239* | 2.6 ± 0.1 | 53.4 ± 3.2 | 53.4 ± 0.8 | 40.9 ± 1.1 | 61.1 ± 0.1 | 18.8 ± 0.0 | 6.3 ± 0.1 | 59.5 ± 4.9 | 8.0 ± 0.1 |
